# Comparative molecular analysis of histopathologic subtypes reveals shared epithelial remodeling yet distinct fibroblast programs in scleroderma lung disease

**DOI:** 10.64898/2026.11.03.763791

**Authors:** Monica M. Yang, Vincent Auyeung, Seon Min Kim, Emily Flynn, Ram P. Naikawadi, Alexey Bazarov, Averey Lea, Fred Deiter, Jessica Neely, Jasleen Kukreja, Michael A. Matthay, Kirk D. Jones, John R. Greenland, Marina Sirota, Paul J. Wolters

## Abstract

**Rationale:** Interstitial lung disease (ILD) drives mortality in systemic sclerosis (SSc), with a subset developing progressive pulmonary fibrosis (PPF) resembling idiopathic pulmonary fibrosis (IPF). Despite recognition of two distinct histopathologic patterns, the cellular basis of nonspecific interstitial pneumonia (NSIP) and usual interstitial pneumonia (UIP) remains poorly understood.

**Objectives:** To define shared and distinct cellular and molecular features of SSc-NSIP and SSc-UIP compared to IPF.

**Methods:** Single-nucleus RNA sequencing and spatial transcriptomics were performed on explanted lungs from patients with SSc-NSIP (n=6), SSc-UIP (n=6), or IPF (n=6), and control donors (n=12) with immunohistochemistry and western blot validation.

**Main Results:** Across 120,254 nuclei from 35 cell types, there was striking similarity across fibrotic subgroups in epithelial, endothelial, and immune compartments, including alveolar type 1 cell loss, pathologic alveolar type 2 (AT2) gene expression, and expansion of COL15A1⁺ endothelial cells and profibrotic macrophages. Diseased AT2s showed activation of the p53/DNA damage pathway. Aberrant basaloid (AB) cells were present in both SSc-ILD subtypes and transcriptionally similar to IPF ABs. They consistently neighbored profibrotic CTHRC1⁺ fibroblasts, even in SSc-NSIP, despite lacking fibroblast foci. Fibroblast heterogeneity differentiated the histopathologic subtypes. Inflammatory fibroblasts were enriched in SSc-NSIP and CTHRC1⁺ fibroblasts in UIP. AB-to-pathologic fibroblast crosstalk was defined by profibrotic signaling, with amplified TGF-β signaling in UIP.

**Conclusions:** Convergent epithelial remodeling is a hallmark of PPF in SSc-ILD and IPF. Fibroblast heterogeneity characterized by altered TGF-β signaling distinguishes SSc-NSIP from SSc-UIP and IPF. Accelerated aging, DNA damage, and p53-dependent senescence pathways may represent shared drivers of PPF.

## Introduction

Interstitial lung disease (ILD) is a common manifestation of systemic sclerosis (SSc) and the leading cause of SSc-related death.^1,2^ Despite earlier recognition and therapeutic advances, a subset of patients with SSc-ILD develop progressive pulmonary fibrosis (PPF) that is fatal without lung transplantation.^3^ Management is limited in part by ineffective classification systems that rely on radiographic and histopathologic patterns rather than molecular phenotyping needed for precision-medicine approaches. SSc-ILD presents in two histopathologic patterns: non-specific interstitial pneumonia (NSIP), characterized by diffuse septal thickening with lymphocytic inflammation, and usual interstitial pneumonia (UIP), characterized by subpleural fibrosis and fibroblast foci adjacent to normal lung.^4,5^ UIP is reported to be less responsive to immunomodulatory therapy and carries a worse prognosis than NSIP.^6^ Despite these differences, little is known about the pathobiology of these subtypes, limiting their clinical utility.

Approximately a quarter of SSc-ILD patients develop PPF.^7^ PPF patients follow a disease course mimicking idiopathic pulmonary fibrosis (IPF), a disease characterized by telomere dysfunction, epithelial cell senescence, and the UIP pattern of fibrosis. PPF in SSc-ILD also shares IPF risk factors, including older age, male sex, and lack of response to immunomodulatory therapy.^8^ The fact that both SSc-NSIP and SSc-UIP can progress to PPF raises the question of whether convergent pathologic cellular and molecular pathways emerge later in the disease course.

Single-cell technologies have advanced the understanding of pathologic cell types in fibrotic lung disease.^9,10^ However, their application to the different histopathologic subtypes of SSc-ILD is limited, with studies primarily focusing on immune and stromal compartments and the SSc-UIP pattern.^11,12^ Here, we provide a detailed cellular and molecular characterization of SSc-ILD with PPF and define shared and distinct aspects of the histopathologic subtypes, with a focus on epithelial cells and their interaction with fibroblasts.

## Methods

### Samples

Archived frozen lung tissues from SSc-ILD and IPF were obtained at transplantation and from unused donors for healthy controls. All lung samples were reviewed by an experienced pathologist to confirm histopathology, and mixed patterns (NSIP/UIP) were excluded from this study. The study was approved by the University of California Institutional Review Board (#13-10738), and all patients gave written consent.

### SnRNA-seq sequencing

Nuclei were isolated from frozen tissue by gentle homogenization with Nuclei Lysis Buffer + RNAse inhibitor, followed by an iodixanol gradient with centrifugation to remove mitochondria and debris. Samples were multiplexed to reduce batch effects, and libraries were generated using the 10x Genomics chemistry v3. Libraries were sequenced on the Illumina NovaSeq 6000, targeting >50,000 reads/cell to characterize rare cell populations. Downstream read alignment and cell calling were conducted using CellRanger.^13^

### Data processing and annotation

Data were analyzed using Seurat.^14^ Ambient RNA was corrected with SoupX, and nuclei with < 500 transcripts or > 3% mitochondrial reads were removed as low-quality cells.^15^ Samples were demultiplexed using Demuxlet, and doublets removed using DoubletFinder.^16,17^ Libraries were integrated using Harmony (Supplementary Figure 1).^18^ Cluster markers were obtained using FindAllMarkers, and, when available, clusters were annotated using published fibrotic lung datasets to avoid introducing new cell names.^19,20^ Nuclei were subsequently subset into epithelial, stromal, and immune compartments, reclustered, and annotated.

### Spatial transcriptomics

Spatial transcriptomics and library generation were performed using the Xenium platform from 10x Genomics with a custom 480-gene lung panel on available FFPE tissues from 17 of 30 individuals. Cells were segmented using the Multimodal Segmentation Kit and the Xenium Onboard Analysis pipeline, and cell annotation was performed by mapping cell labels from the single-nucleus dataset to ensure concordant cell labeling.

### Downstream analysis and statistics

Cell type proportions were compared between groups using the Wilcoxon rank-sum test and Kruskal-Wallis test. Differential expression was performed on sample-level pseudobulked counts using EdgeR (padj <0.05 and log2FC >1).^21^ PCA distances were compared using centroid distances and DEG correlation was calculated using the Spearman coefficient. Pathway analysis was performed with ClusterProfiler, and previously published inflammation, ECM, and senescence module scores were leveraged.^22–25^ Epithelial trajectories were inferred using Monocle3, and cell-cell communication was assessed using CellChat (p<0.05).^26^ Cell neighborhood analysis, the spatial coordinates of cell centroids, defined by the Xenium instrument on-board analysis (10X Genomics), were organized into a K-D tree, and each cell’s fixed-radius nearest neighbors were identified using dbscan.^27^

## Results

### Cohort Characteristics

Lung tissues from 30 individuals, 12 SSc-ILD (6 SSc-NSIP, 6 SSc-UIP), 6 IPF, and 12 healthy unused donors (HC), were included (Figure 1A-B, Supplementary Table 1). The mean age of the cohort was 53.8 ± 9.7 years and similar across the four groups (p=0.22). The cohort was 63% female (p=0.83) and predominantly white (p=0.76).

**Figure 1.**
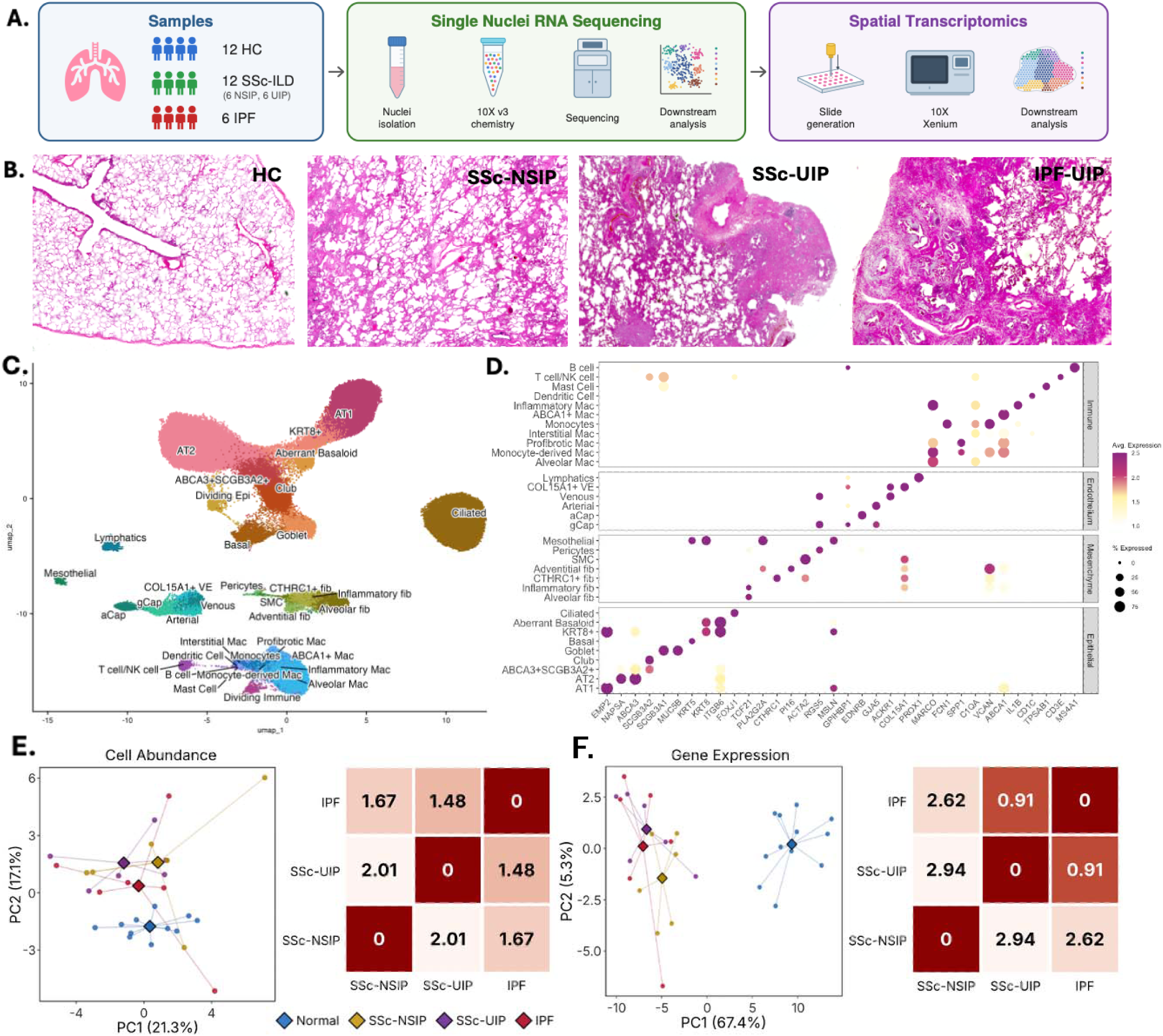
Comparative molecular analysis reveals shared and distinct features among SSc-ILD subtypes and IPF. A) Study overview, including lung tissue from 12 healthy controls, 6 SSc-NSIP, 6 SSc-UIP, and 6 IPF. B) Representative images of H&E with all tissues reviewed by an expert pathologist. C) UMAP of snRNAseq with 120,254 cells. D) Dotplot of canonical markers of 35 cell clusters. E-F) PCA and centroid distances by E) Cell Abundance and F) Gene Expression with the diamond representing the group centroid.

There were more former and current smokers in IPF (83%) compared to SSc-ILD (16%) or HC (41%) (p = 0.04).

### Single-nuclei RNAseq captures diverse cell types

The final dataset included 120,254 nuclei and 33,706 unique genes, averaging 2723 genes and 4274 transcripts (UMIs) per cell. Thirty-five cell types were identified (Figure 1C-D), including pathologic epithelial cells previously poorly characterized in SSc-ILD, such as Aberrant Basaloid (ABs), KRT8+, and alveolar type 2 (AT2) club transitional cell (labeled ABCA3^+^SCGB3A2^+^).

At the whole-lung level, principal component analysis (PCA) demonstrated clear separation between normal and fibrotic lungs in cell composition and gene expression (Figure 1E-F) driven by alveolar type 1 cells (AT1) in HC, and pathologic epithelial cells and fibroblasts (CTHRC1^+^, Inflammatory) in fibrotic lungs (Supplementary Figure 2). The three fibrotic subgroups had similar cell composition. Transcriptionally, however, SSc-UIP was closer to IPF compared to SSc-NSIP, with a threefold smaller centroid distance. This suggests that while cellular composition is similar between fibrotic subtypes, the UIP pattern may share a common gene expression profile.

### Profibrotic Macrophages are shared among IPF and SSc-ILD

The immune compartment included 10,946 nuclei and 12 clusters (Figure 2A-B). Macrophages were most common, followed by T/NK cells, monocytes, dendritic cells, mast cells, and B cells. Distinct macrophage subsets included alveolar, interstitial, inflammatory, cholesterol-laden (ABCA1^+^), monocyte-derived, and profibrotic. Across fibrotic subgroups, there was a loss of alveolar macrophages (p=0.17), with reciprocal increases in profibrotic macrophages and dividing immune cells compared to HC (p < 0.01, Figure 2C, Supplementary Figure 3). Immune composition was similar across the fibrotic groups, with SSc-UIP and IPF having the closest PCA centroids (Figure 2D).

**Figure 2.**
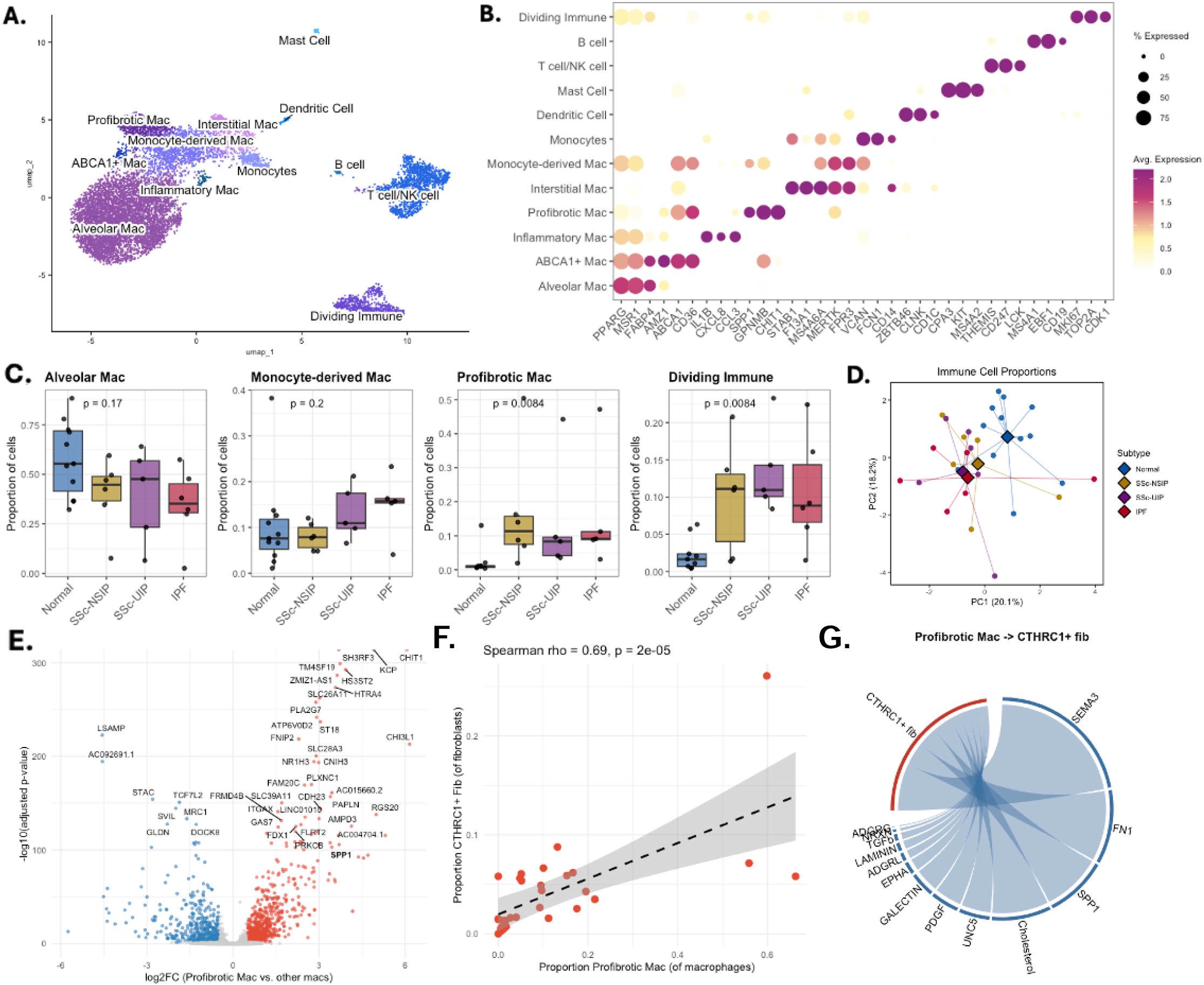
Profibrotic Macrophages are expanded in SSc-ILD and IPF. A) UMAP of immune cells. B) DotPlot of canonical markers of 12 clusters. C) Cell proportions of select immune subsets by Kruskal-Wallis. D) PCA of immune cell abundance. E) Volcano plot of DEGs in profibrotic macrophages vs. all other macrophages across all samples. F) Across IPF and SSc-ILD, the proportion of profibrotic macs correlates with CTHRC1+ fibs. G) Chord diagram of signaling from profibrotic macs to CTHRC1+ fibs in fibrotic lungs.

Profibrotic macrophages, hypothesized to arise from monocyte-derived macrophages, demonstrated upregulation of fibrosis– and inflammation-related markers (CHIT1, SPP1, PLA2G7, and CHIL31) with loss of canonical macrophage markers (MRC1 and STAC) (Figure 2E) across all fibrotic subgroups.^28^ In IPF, these macrophages are proposed to interact with pathologic fibroblasts and promote fibrosis.^29^ Consistent with this, profibrotic macrophage abundance was associated with CTHRC1^+^ fibroblast abundance, and inversely related to alveolar macrophage abundance (Figure 2F, Supplementary Figure 4). This was recapitulated in the spatial data, where neighborhood analysis revealed that, among fibroblasts, CTHRC1^+^ fibroblasts were among the most common neighboring subsets to profibrotic macrophages across ILD subtypes. Cell communication analysis between profibrotic macrophages and CTHRC1^+^ fibroblasts identified profibrotic pathways (FN1, SPP1, PDGF, TGF-β) and adhesion and migration signaling pathways (SEMA3 and GALECTIN) (Figure 2G).

### Endothelial changes are shared across fibrotic lung diseases, while fibroblast subtypes distinguish NSIP and UIP

The stromal compartment consisted of 13 clusters, including 4 fibroblast and 6 endothelial cells (Figure 3A-B). Endothelial cell types included arterial, venous, aerocyte capillary (aCap), general capillary (gCap), lymphatics, and a COL15A1^+^ venous endothelial (VE) cluster.^10^ Cell abundance analysis revealed a trend toward loss of aCaps across fibrotic subgroups, with an enrichment in the COL15A1^+^ VE in both SSc-ILD and IPF (p=0.07, p<0.01, Figure 3C-D), and gene expression analysis revealed shared pathologic gene expression across fibrotic lung diseases (Supplementary Figure 5). The COL15A1^+^ VE showed a loss of canonical endothelial markers and a reciprocal increase in genes associated with endothelial-mesenchymal transition (EndoMT) (Figure 3E), with enrichment of pathways related to ECM interactions, integrin signaling, and cellular senescence (Figure 3F). Spatially, the COL15A1^+^ VE neighborhood was similar in IPF and SSc-ILD, comprising lymphocytes, other VEs, and inflammatory and CTHRC1^+^ fibroblasts (Figure 3G).

**Figure 3.**
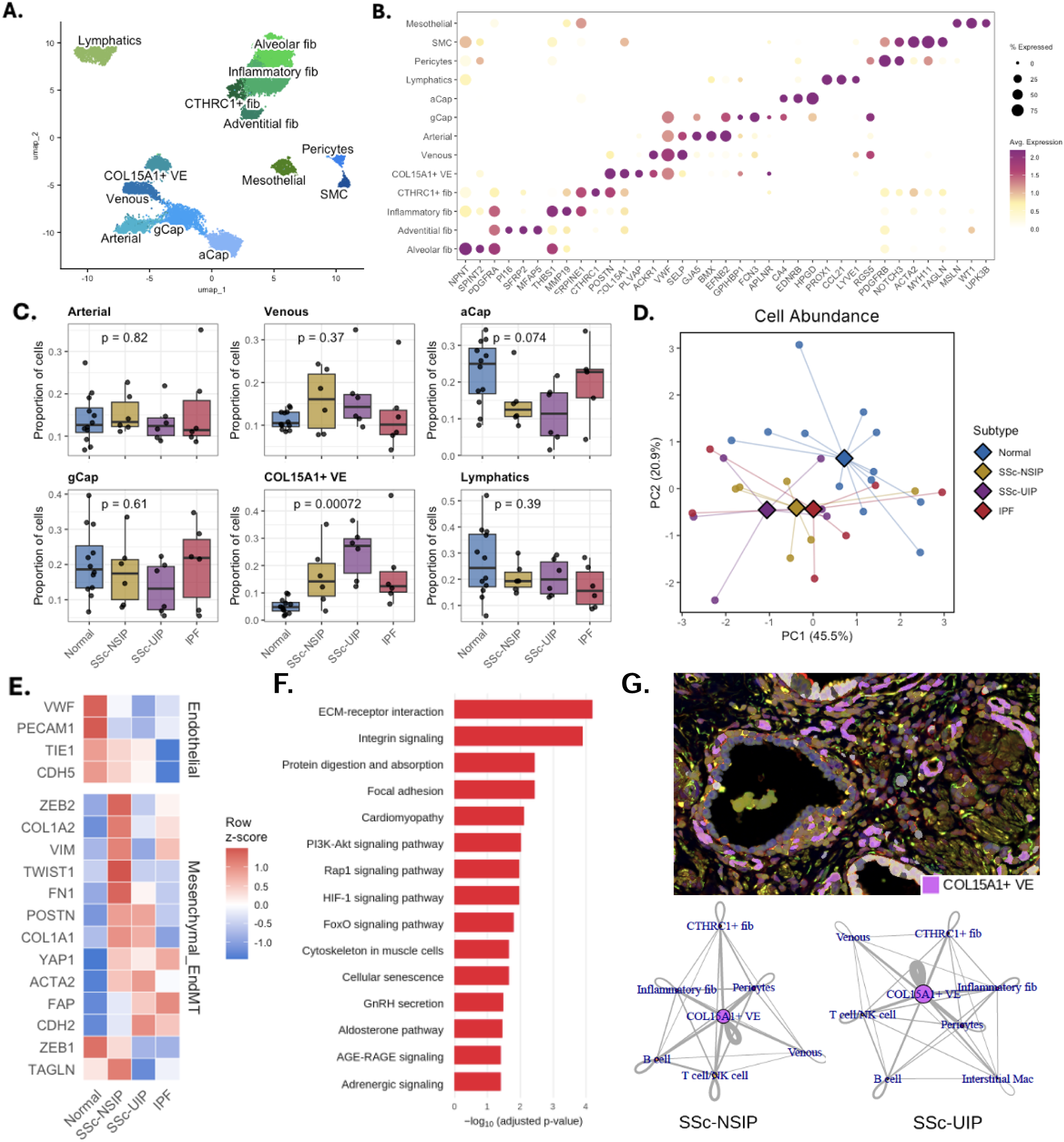
COL15A1+ VE are expanded in SSC-ILD and IPF and characterized by EndoMT features. A) UMAP of endothelial cells. B) DotPlot of canonical markers of 13 clusters. C) Cell proportions of endothelial subsets by Kruskal-Wallis. D) PCA of endothelial cell abundance. E) Heatmap of endothelial and EndoMT genes. F) Hallmark pathways enriched in COL15A1+ VE across diseases. G) Xenium image of COL15A1+ cells with neighborhood analysis of SSc-NSIP and SSc-UIP.

Fibroblast cell types included alveolar, adventitial, inflammatory, and profibrotic (CTHRC1^+^) fibroblasts.^20^ Cell abundances differed significantly between normal and fibrotic subgroups, with greater similarity among UIP fibrosis patterns, a pattern also observed when comparing DEGs (Figure 4A-C). There was loss of alveolar fibroblasts across all fibrotic subgroups, with enrichment of inflammatory fibroblasts, greatest in SSc-NSIP, and CTHRC1^+^ fibroblasts, greater in SSc-UIP and IPF. Importantly, both pathologic fibroblast subtypes were present in each fibrosis pattern. Inflammatory fibroblasts were enriched in inflammation-driven modules, including TNF-alpha, interferon, and IL6, while the CTHRC1^+^ fibroblasts showed increased profibrotic modules, including the core fibrosis module, collagens, and ECM regulators (Figure 4D). Both pathologic fibroblasts showed higher senescence module scores than alveolar and adventitial fibroblasts, with CTHRC1^+^ having the highest score. At the individual gene level, SSc-NSIP inflammatory fibroblasts showed stronger expression of inflammatory markers, whereas IPF inflammatory fibroblasts showed higher expression of profibrotic genes. Similarly, in CTHRC1^+^ fibroblasts, expression of profibrotic markers was stronger in the UIP pattern of fibrosis (Supplementary Figure 6).

**Figure 4.**
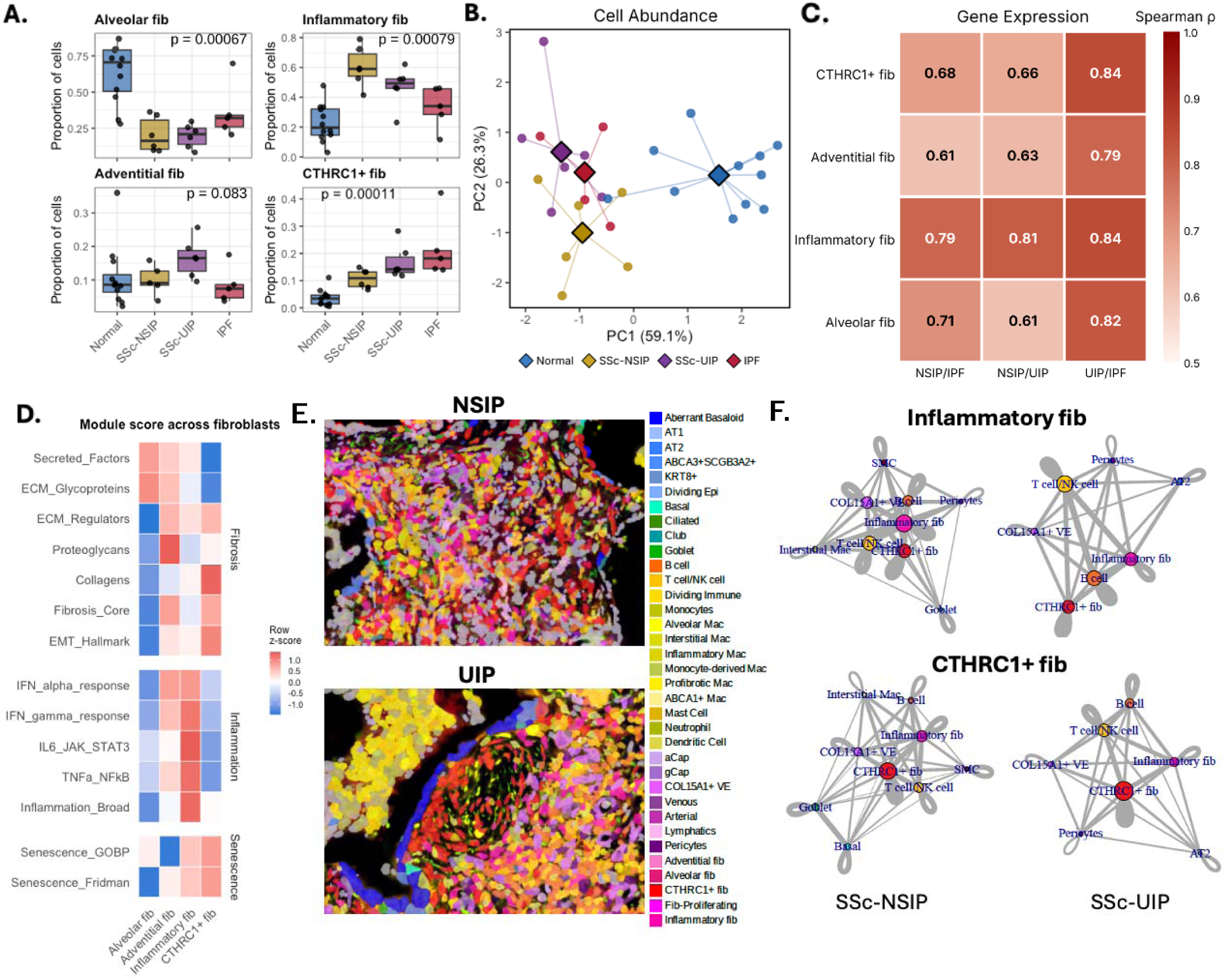
Distinct fibroblast subtypes and spatial niches arise in SSc-NSIP vs. SSc-UIP. A) Cell proportions of fibroblast subsets by Kruskal-Wallis. B) PCA of fibroblast abundance. C) Spearman correlation of disease vs. normal log fold changes between pairs of subtypes by fibroblast subset. D) Fibrosis, inflammation, and senescence module scores across fibroblast subsets. E) Xenium images of inflammatory and CTHRC1+ fibroblast niches and F) spatial neighborhood analysis in SSc-NSIP and SSc-UIP.

In addition to their differences in cell proportion, there were striking differences in the spatial architecture of pathologic fibroblasts between subtypes. Neighborhood analysis revealed that inflammatory fibroblasts colocalized primarily with immune cells, COL15A1^+^ VE, and CTHRC1^+^ fibroblasts in both SSc-UIP and SSc-NSIP. However, compared to UIP, when inflammatory fibroblasts were present in NSIP, there was increased surrounding density (number and proximity) of other pathologic fibroblasts, lymphocytes, and pathologic endothelial cells, potentially corresponding to areas of inflammatory infiltrates (Figure 4E). CTHRC1^+^ fibroblasts have been described as localizing to fibroblast foci, a key pathologic finding in UIP.^30^ In this dataset, CTHRC1^+^ fibroblasts were present in both NSIP and UIP, and spatially, while the cell distances varied, their neighborhood composition was surprisingly similar between the two patterns (Figure 4F). In SSc-UIP, CTHRC1^+^ fibroblasts were associated with fibroblast foci, whereas in SSc-NSIP, CTHRC1^+^ fibroblasts appeared more dispersed in the dense cellular infiltrate. Interestingly, both pathologic fibroblasts in SSc-ILD (NSIP and UIP) were in closer proximity to the COL15A1^+^ pathologic VE than in IPF, emphasizing the role of endothelial dysfunction in the disease.

### Epithelial cell remodeling demonstrates shared patterns across fibrotic lung disease and overlapping pathologic gene expression

Given that epithelial remodeling is central to IPF yet understudied in SSc-ILD, we focused our in-depth analysis on the epithelial compartment. Re-clustering revealed nine cell types, including six normal lung epithelial cell types: alveolar type 2 (AT2), AT1, club, goblet, basal, and ciliated cells, and three disease-associated: AB, KRT8^+^, and ABCA3^+^SGB3A2^+^ cells (Figure 5A). All fibrotic lung diseases showed a profound loss of AT1s (p=0.003), with a reciprocal increase in airway cells, including club and goblet cells (p=0.01, p=0.01, Figure 5B). PC distances for cell abundance and gene expression mirrored the PCA results from the whole-lung dataset (Figure 5C). Among canonical epithelial cells, DEGs showed the strongest correlation between SSc-UIP and IPF, whereas pathologic cells showed strong gene correlation across all fibrotic subtypes (Figure 5D). Although AT2s were not depleted in fibrotic disease, their gene expression was significantly changed compared to HCs (Figure 5E). Diseased AT2s showed loss of canonical AT2 markers (SFTPC, SFTPA1, LAMP3), with the greatest loss in IPF, followed by SSc-UIP, and then SSc-NSIP. The p53 pathway was the most highly enriched pathway in diseased AT2 cells, along with other pertinent pathways, including UV Response (DNA Damage), ROS, and EMT (Figure 5F).

**Figure 5.**
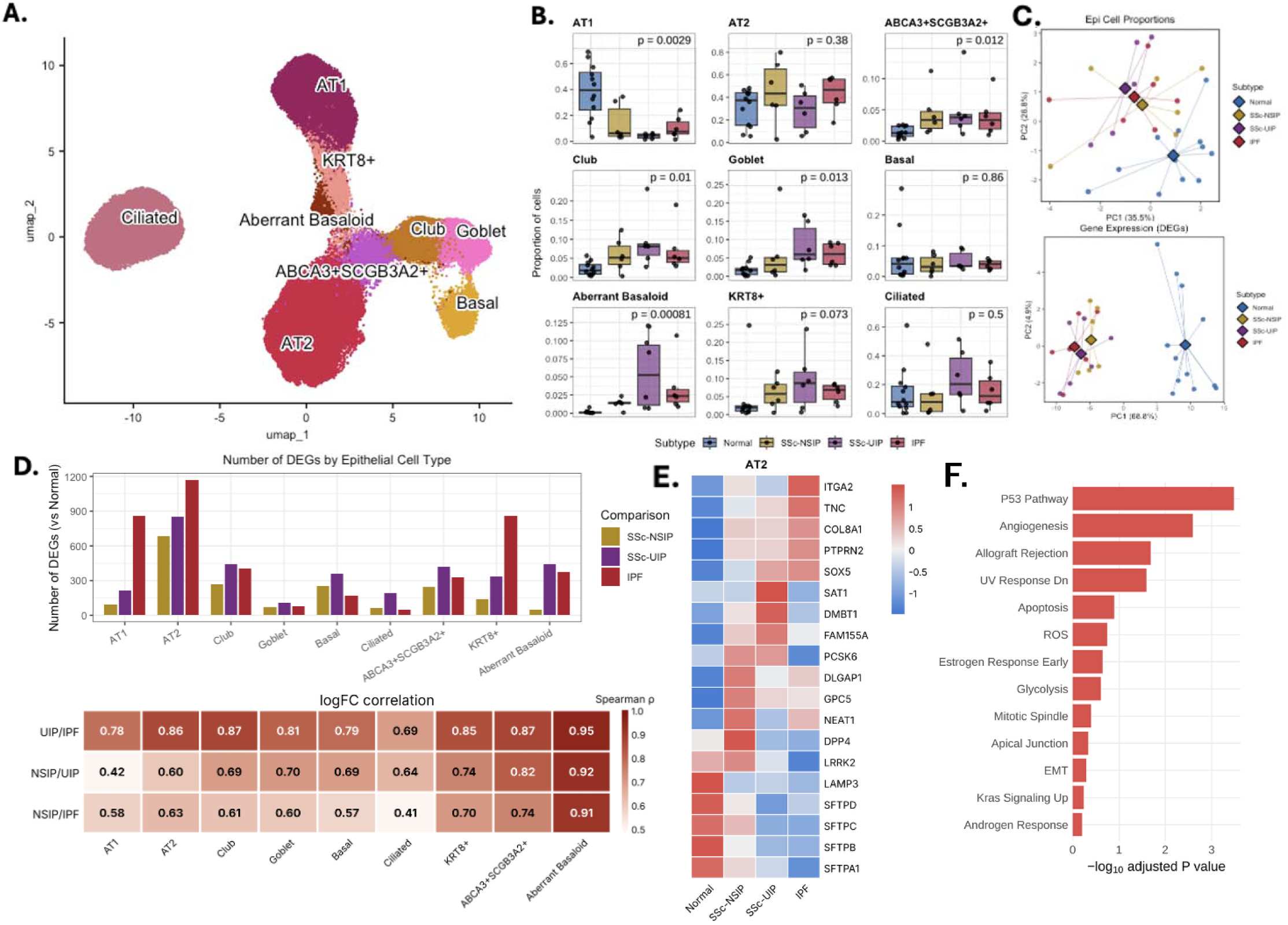
Epithelial remodeling in SSc-ILD and IPF shows loss of AT1 and emergence of pathologic AT2. A) UMAP of epithelial cells. B) Cell proportions of epithelial subsets by Kruskal-Wallis. C) PCA of epithelial cell abundance and gene expression. D) Number of DEGs by epithelial cell and their Spearman correlation between pairs of disease subtypes. E) Heatmap of selected AT2 DEGs. F) Hallmark pathways enriched in AT2 DEGs across fibrotic lung diseases.

### Emergence of pathologic and transitional cell types characterized by aging and senescence pathways

Three pathologic epithelial cell types were identified in our dataset. Two transitional cell clusters were present, including an AT2-club hybrid cell (ABCA3^+^SGB3A2^+^), which resembled previously described hybrid clusters (e.g., SFTPC^+^SCGB3A2^+^, AT0), and a KRT8^+^ cell, which has been described as a transitional cell state that arises in the context of lung injury and has been found in IPF, ARDS, and COVID.^31–33^ KRT8^+^ cells were increased across the disease spectrum and expressed markers like ABs. We captured a robust population of ABs enriched in SSc-ILD and IPF, with a preferential abundance in the UIP pattern.

Aberrant basaloid cells were characterized by markers of cellular senescence, extracellular matrix, and collagen-related genes, and had the highest ECM and senescence module scores of epithelial cell types (Figure 6A). EMT was the most highly enriched pathway, followed by TGF-β signaling (Figure 6B). Other relevant pathways included inflammatory signaling (TNF-alpha, IFN-gamma, and inflammatory response) and aging and senescence, including the UV response (DNA damage) and the p53 pathway. Spatially, ABs were consistently near CTHRC1^+^ fibroblasts across all subtypes, despite differences in AB cell proportions between subtypes, and the lack of fibroblast foci in SSc-NSIP (Figure 6C). When present, the number of neighboring CTHRC1^+^ fibroblasts was very similar (Figure 6D). This conserved spatial relationship suggests their cell interaction may be relevant to the PPF disease phenotype. In IPF, AT2 cells have been proposed to give rise to ABs.^34^ In our dataset, pseudotime analysis suggests that ABs emerge along the AT2-to-AT1 differentiation path (Supplementary Figure 7). Reclustering cells along this path revealed two trajectories: one from AT2 to AT1 and another from AT2 to AB, both passing through a KRT8 intermediate (Figure 6E). Comparing gene expression changes over these trajectories in fibrotic lungs, as well as that of AT2 to AT1 in HCs, there was a gain of markers of cellular senescence (GDF15, CDKN1A, CDKN2A, SERPINE1) and fibrosis (COL1A1, COL7A1) along the AB arm (Figure 6F).

**Figure 6.**
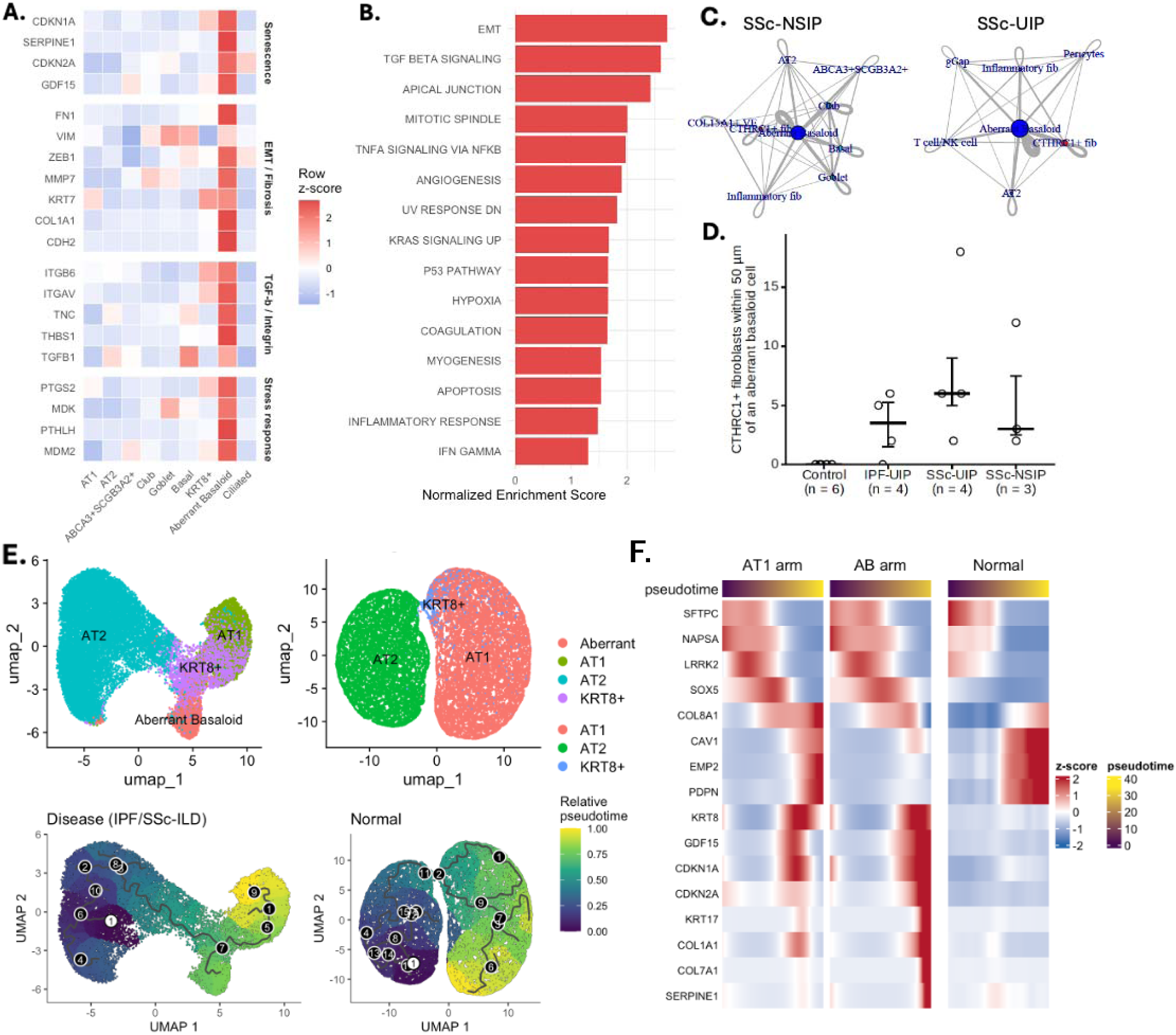
Aberrant Basaloid cells show senescence and EMT programs and may emerge from pathologic AT2. A) Heatmap of pathologic gene expression in ABs. B) Hallmark pathway enrichment in ABs. C) Spatial neighborhood analysis around ABs in NSIP vs. UIP. D) Number of CTHRC1+ fibroblasts within 50um of ABs by subtype. E) Pseudotime analysis of AT2 to AT1 differentiation in disease vs. normal lungs. F) Gene expression changes along pseudotime comparing AT2 to AT1 vs. AT2 to AB.

To validate transcriptomic signatures of senescence and DNA damage, protein expression levels of relevant markers in whole-lung lysates were assessed (Figure 7A-B). Immunoblot of whole-lung lysates showed markedly increased levels of phosphorylated CHK1, CHK2, and p53 across fibrotic lung diseases, implicating activation of the DNA damage response (DDR) across disease subtypes. The senescence marker p16 and the TGF-β signaling marker pSMAD3 were also increased in SSc-ILD and IPF, mirroring transcriptomic findings.

**Figure 7.**
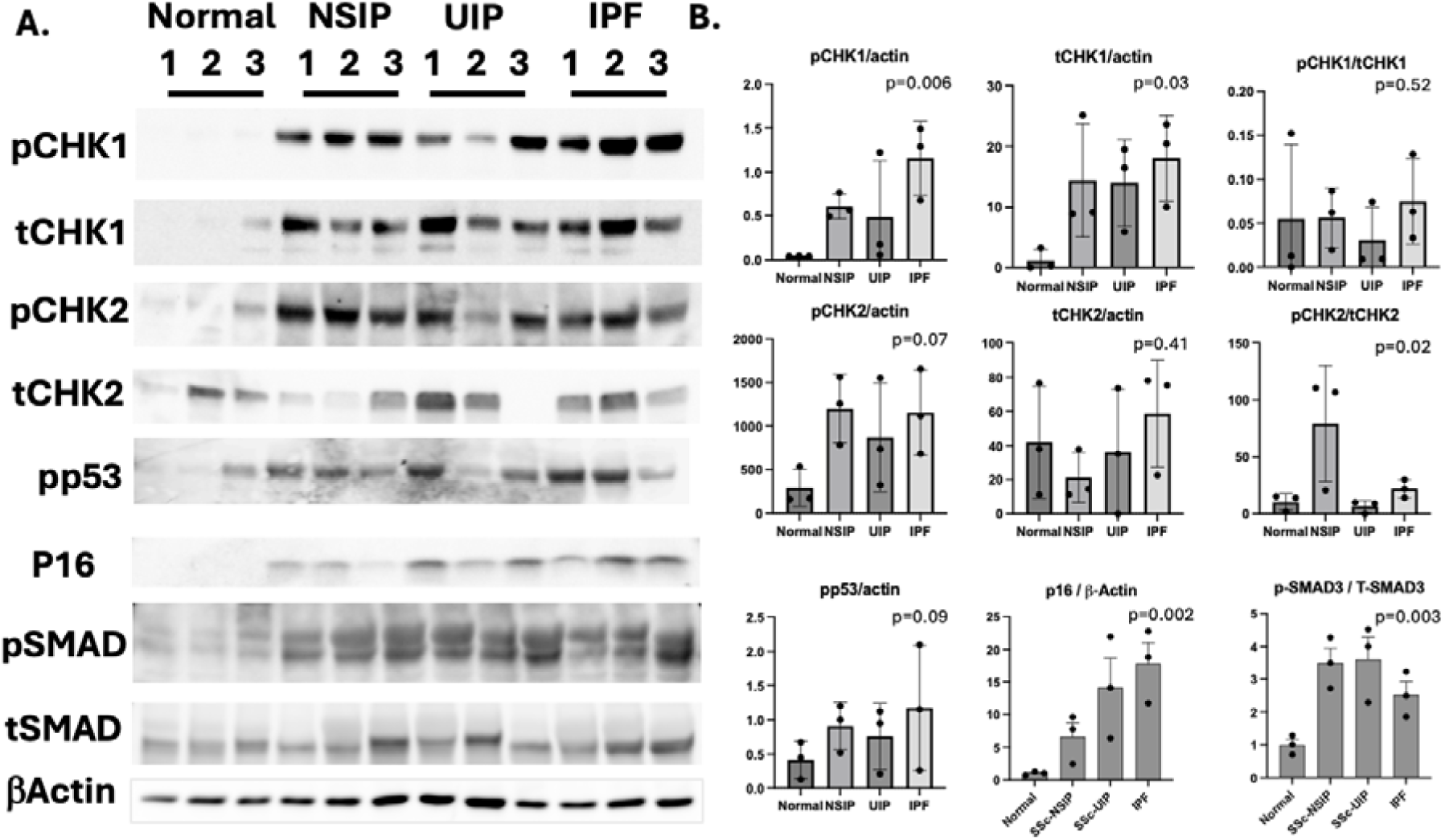
Activation of DNA damage, senescence and TGF-β in SSc-ILD and IPF. A) Western blots of pCHK1, tCHK1, pCHK2, tCHK2, phospho-p53, p16, pSMAD3, and tSMAD3 in lung lysates from normal, SSc-NSIP, SSc-UIP, and IPF (n = 3 per group). β-actin is the loading control. B) Densitometric quantification. Bars show mean ± [SD/SEM]; dots are individual samples.

### Pathologic epithelial and fibroblast cells participate in profibrotic crosstalk

Cell communication analysis revealed that most predicted interactions occurred between epithelial cells and fibroblasts (Supplementary Figure 9). ABs, KRT8+, and four fibroblast subtypes had the greatest crosstalk, so subsequent analysis focused on these cell interactions. Compared to HCs, differential cell interactions were increased among diseases, with SSc-UIP and IPF showing more numerous and stronger interactions than SSc-NSIP (Figure 8A). In SSc-ILD, AB-to-fibroblast signaling was skewed towards pathologic fibroblasts, and away from alveolar and adventitial fibroblasts (Figure 8B). The shift favored CTHRC1+ in SSc-UIP and inflammatory fibroblasts in SSc-NSIP. Importantly, these analyses were independent of cell number, reinforcing the cell abundance and spatial neighborhood findings.

**Figure 8.**
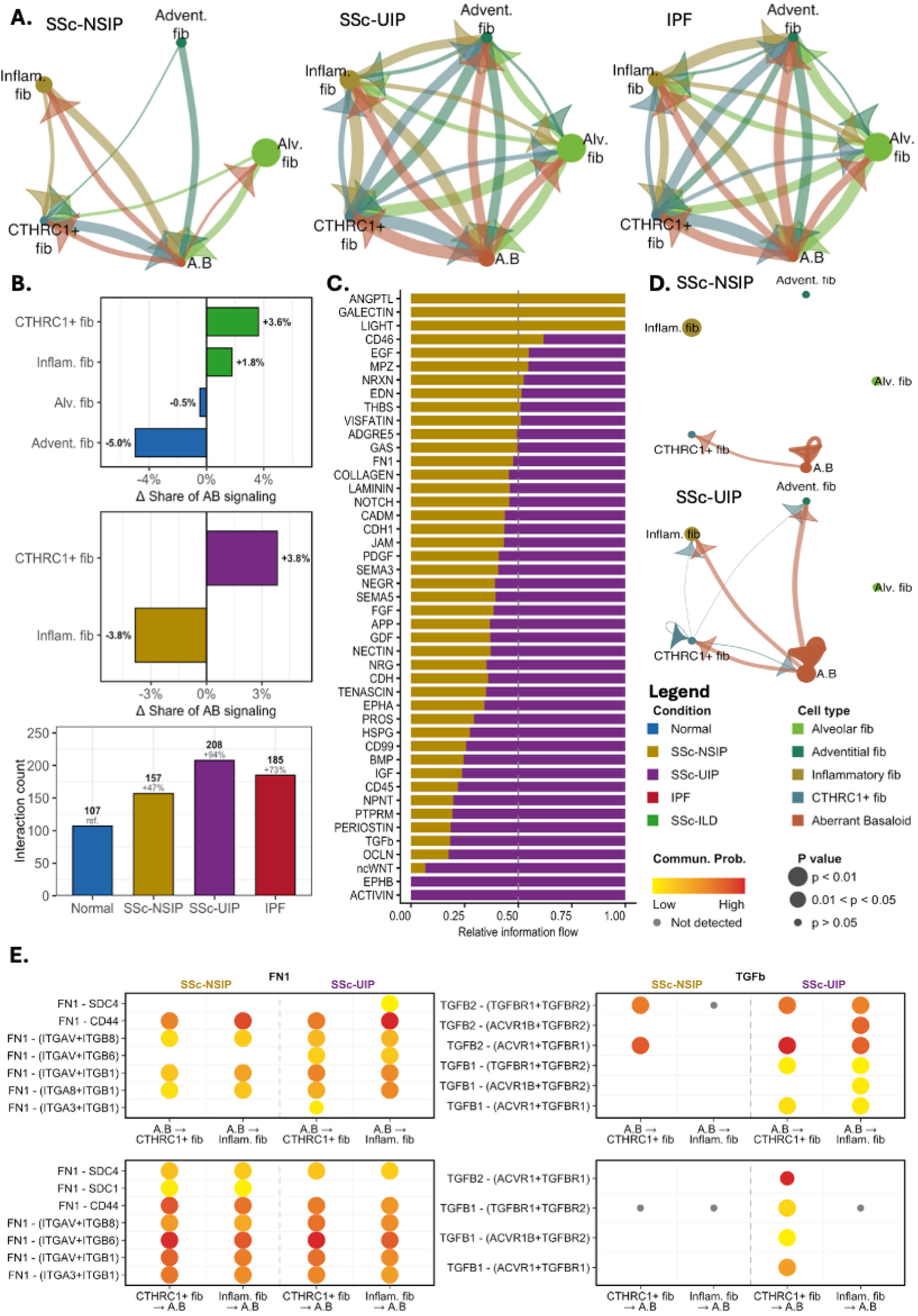
Aberrant basaloid and pathologic fibroblasts participate in profibrotic signaling in SSc-ILD and IPF. (A) Circle plots of predicted interactions between aberrant basaloid (AB) cells and fibroblast subsets in SSc-NSIP, SSc-UIP and IPF. (B) Change in each fibroblast subset’s share of AB signaling, and total interaction counts per group. (C) Relative information flow of signaling pathways in S c-NSIP vs. SSc-UIP. (D) TGF-β signaling between AB cells and fibroblast subsets in SSc-NSIP (top) vs. SSc-UIP (bottom). (E) Dot plots of FN1 and TGF-β ligand–receptor pairs between AB cells and each pathologic fibroblast.

Epithelial-fibroblast communication differed further between SSc-ILD subtypes. SSc-NSIP was preferentially enriched for pathways relevant to inflammatory remodeling (ANGPTL, GALECTIN, LIGHT, CD46) (Figure 8C), and SSc-UIP was preferentially enriched for profibrotic signaling (ACTIVIN, PERIOSTIN, non-canonical WNT signaling) (Figure 8C). Notably, TGF-β was among the pathways most skewed toward SSc-UIP. In SSc-NSIP, TGF-β signaling was largely restricted to a CTHRC1+ fibroblast-AB interaction, whereas in SSc-UIP, ABs engaged in robust TGF-β signaling to CTHRC1^+^, inflammatory, and adventitial fibroblasts as well (Figure 8D). Ligand-receptor analysis demonstrated bidirectional TGFB1/TGFB2 signaling between ABs and pathologic fibroblasts through multiple receptor complexes in SSc-UIP (Figure 8E). By contrast, FN1 and Collagen signaling were robust between ABs and fibroblasts in both SSc-ILD subtypes, indicating that while global profibrotic signaling is a shared feature of SSc-ILD, TGF-β signaling is preferentially amplified in the fibrotic UIP pattern of fibrosis.

## Discussion

This study provides an in-depth molecular characterization of the major histopathologic subtypes of SSc-ILD with a PPF phenotype. By comparing to IPF, the analysis investigated whether each subtype’s molecular profiles are driven more by the underlying disease or histopathologic subtype. This dataset enables robust characterization not only of distinct histopathologic subtypes but also pathologic cell types previously under-characterized in SSc-ILD. Although there were unique compartment– and cell-type-specific findings between the fibrotic subtypes, the overall finding was a striking similarity among SSc-NSIP, SSc-UIP, and IPF, suggesting common pathobiology across fibrotic lung diseases with the PPF phenotype. In particular, aging-associated pathways were upregulated across fibrotic lung diseases. The histopathologic subtypes appear to be primarily defined by differences in fibroblast architecture, with the UIP pattern driven by enhanced TGF-β signaling.

Given the differences in NSIP and UIP histopathology, it was surprising to find that the predominant pathobiologic changes in SSc-NSIP and SSc-UIP overlapped with those in IPF. Most notable were shared elements of epithelial remodeling, including a loss of AT1s, an increase in airway cells, and the emergence of KRT8^+^ and ABs. ABs, which are well described in IPF, have been reported in SSc-ILD, but in very limited numbers and not in the NSIP pattern of fibrosis.^12,35^ Here, ABs are present in a similar proportion to IPF in both SSc-NSIP and SSc-UIP and are transcriptionally identical across diseases. Spatially, to date, these cells have primarily been associated with fibroblast foci, however, we show ABs and their spatial relationship to CTHRC1^+^ fibroblasts are conserved in SSc-NSIP and present within the dense cellular infiltrate characteristic of its histopathology. Given their presence across fibrotic lung diseases, ABs and their associated remodeling may be shared features across PPF subtypes, extending beyond UIP fibroblast foci and reinforcing their role in lung fibrosis progression.

Interestingly, the loss of AT2 cells observed in scRNA-seq datasets was less prominent in this snRNA-seq analysis.^9,10^ However, AT2 cells in fibrotic lungs had the greatest deviation in gene expression relative to HCs, indicating a diseased phenotype despite similar numbers. It is possible that fragile, diseased, AT2 cells may be lost during single-cell processing, explaining their underrepresentation in single-cell datasets. Overall, AT2s in SSc-ILD and IPF lost expression of canonical AT2 cell markers and gained expression of profibrotic and proinflammatory genes. Importantly, changes were more prominent in UIP than in NSIP lungs, suggesting IPF and SSc-UIP AT2s are transcriptionally more similar.

Senescence reprogramming of AT2s has been shown to drive IPF progression and appears similarly relevant in SSc-ILD.^36,37^ The p53 and DNA damage pathways were the most differentially expressed pathways in diseased AT2s, and associated proteins were increased in lung lysates of SSc-ILD and IPF. Trajectory analysis suggests that senescent AT2s transition towards ABs rather than fully differentiating into AT1s, resulting in a stalled pathologic state. Prior reports describe the relevance of aging and senescence to SSc-ILD, including telomere attrition in AT2 cells, shortened peripheral blood leukocyte telomere length associated with SSc-ILD progression, and upregulation of the p53 pathway in bulk RNA-seq data.^38,39^ These data extend these findings by showing p53 activation in AT2s and AB, highlighting p53 and the DNA damage response as possible mechanistic targets in PPF.

Across fibrotic lung diseases, shared biology includes endothelial cell changes and the emergence of pathologic endothelial cell subsets. Although SSc is characterized by endothelial dysfunction, the role of endothelial dysfunction is poorly defined in SSc lung disease and IPF. This study found a loss of aCap cells, a finding also associated with aging lungs.^40^ This loss mirrors the loss of AT1 cells and reinforces impaired gas exchange in the fibrotic lung. Furthermore, all fibrotic lungs had expansion of COL15A1^+^ endothelial cells, which colocalize with CTHRC1^+^ fibroblasts and express EndoMT-associated genes. The combined loss of angiogenic gene expression and emergence of profibrotic endothelial gene expression highlight a shared role for endothelial cell dysfunction in progressive SSc-ILD and IPF.

The greatest differences between NSIP and UIP emerged in fibroblast subsets. Single-cell studies have advanced the understanding of fibroblast heterogeneity, including the identification of inflammatory and profibrotic fibroblast subsets.^20,41^ Notably, the original dataset included three SSc-ILD lungs, which, on further review, were all NSIP pattern, likely explaining the predominance of inflammatory fibroblasts in SSc-ILD in that report. Consistent with these findings, this comparative analysis of SSc-NSIP vs. SSc-UIP shows that proinflammatory gene modules and immunomodulatory cell communication pathways are preferentially active in SSc-NSIP, supporting the idea that NSIP may, in part, be a more immunologically driven fibrogenic process than UIP. Conversely, CTHRC1^+^ fibroblasts were more abundant in UIP across diseases, consistent with their dominant role as matrix-producing fibroblasts enriched in regions of active fibrogenesis, such as fibroblast foci. Spatial and cell communication analyses place these cells near ABs, where they preferentially engage in profibrotic signaling, most notably through TGF-β. The relative lack of inflammatory signaling and heightened profibrotic signaling in SSc-UIP may, in part, explain its limited response to immunomodulatory therapies.

Collectively, these data suggest that epithelial and stromal compartments drive shared disease features across SSc-NSIP, SSc-UIP, and IPF. We propose that pathologic epithelial remodeling underlies the progressive phenotype common to a subset of SSc-ILD and IPF. PPF may be driven in part by aging and senescence-associated AT2 cell reprogramming, resulting in AT2 cells that are trapped in a transitional state, leading to the loss of AT1 cells. These senescent AT2 cells may also give rise to ABs, which propagate lung fibrosis. The remodeling caused by epithelial dysfunction is likely exacerbated by endothelial dysfunction in both SSc-ILD and IPF. With epithelial dysfunction underpinning PPF, the histopathologic subtypes may be driven by fibroblast subtypes: TGF-β-mediated CTHRC1^+^ fibroblasts drive the UIP pattern, whereas SSc-NSIP is enriched for inflammatory fibroblasts. This framework also offers a mechanistic basis for the reported transition from NSIP to UIP, whereby inflammatory fibroblasts convert to CTHRC1^+^ fibroblasts via TGF-β signaling.

This study is the first in-depth characterization of SSC-ILD histopathologic subtypes. The depth of sequencing enabled a granular understanding of healthy and pathological cells. Although archived lung samples enabled controlling for factors such as age, sex, and race, human biology is inherently heterogeneous, as reflected in the data. Single-nucleus RNA-seq enables profiling of fragile cell types but restricts analysis to intranuclear RNA, reducing transcript counts and limiting analysis of low-RNA cell types. This particularly impacted immune cells, which appear to be underrepresented in the dataset. Because scRNAseq appears superior for characterizing immune cells, these two approaches are complementary. Additionally, droplet-based methods were used, which increased ambient RNA. While removed computationally, this resulted in the exclusion of cells from analysis.

In conclusion, multimodal profiling of SSc-ILD histopathologic subtypes and IPF reveals a convergent molecular framework in which the clinical PPF phenotype may arise from shared epithelial and endothelial cell remodeling, with divergent fibroblast biology determining the histopathologic subtype. These findings implicate aberrant aging and senescence programs, particularly DNA damage and p53 pathways, as shared drivers of epithelial dysfunction that promote the development of aberrant basaloid cells, which in turn promote fibrotic remodeling through pathologic epithelial-fibroblast crosstalk. This crosstalk differs between SSc-NSIP and SSc-UIP, with the latter characterized by robust TGF-β signaling. Overall, these findings suggest that, despite distinct etiologies and histopathology, progressive ILDs have more biological similarities than differences.

## Acknowledgment

This work was supported by grants from the Scleroderma Research Foundation, the Rheumatology Research Foundation, Nina Ireland Program for Lung Health, and NIH R01 HL-139897.

## Data Availabilit

Data will be made available through the Gene Expression Omnibus at the time of publication.

