## Supplemental Tables/Figures for "Comparative molecular analysis of histopathologic subtypes reveals shared epithelial remodeling yet distinct fibroblast programs in scleroderma lung disease"

**Supplementary Table 1: Subject Demographics**

| Characteristic | HC (n=12) | SSc-NSIP (n=6) | SSc-UIP (n=6) | IPF (n=6) | p-value |
| --- | --- | --- | --- | --- | --- |
| **Age, years** |  |  |  |  |  |
| Mean ± SD | 48.1 ± 18.6 | 50.7 ± 10.7 | 52.2 ± 10.6 | 62.3 ± 5.0 | 0.22 |
| Median [IQR] | 50.5 [36.3–60.5] | 51.5 [42.8–57.3] | 49.0 [45.3–61.8] | 62.0 [59.3–64.8] |  |
| Range | 19–78 | 37–65 | 40–65 | 56–70 |  |
| **Sex, n (%)** |  |  |  |  | 0.83 |
| Female | 7 (58.3) | 4 (66.7) | 5 (83.3) | 3 (50.0) |  |
| Male | 5 (41.7) | 2 (33.3) | 1 (16.7) | 3 (50.0) |  |
| **Race/ethnicity, n (%)** |  |  |  |  | 0.76 |
| White | 6 (50.0) | 1 (16.7) | 4 (66.7) | 5 (83.3) |  |
| Black | 1 (8.3) | 1 (16.7) | 1 (16.7) | 1 (16.7) |  |
| Hispanic | 3 (25.0) | 3 (50.0) | 1 (16.7) | 0 (0.0) |  |
| Asian | 1 (8.3) | 1 (16.7) | 0 (0.0) | 0 (0.0) |  |
| American Indian | 1 (8.3) | 0 (0.0) | 0 (0.0) | 0 (0.0) |  |
| **Smoking, n (%)** |  |  |  |  | 0.04 |
| Never | 6 (50.0) | 5 (83.3) | 5 (83.3) | 1 (16.7) |  |
| Former | 3 (25.0) | 1 (16.7) | 0 (0.0) | 5 (83.3) |  |
| Current | 2 (16.7) | 0 (0.0) | 1 (16.7) | 0 (0.0) |  |
| Unknown | 1 (8.3) | 0 (0.0) | 0 (0.0) | 0 (0.0) |  |

SSc, systemic sclerosis; NSIP, nonspecific interstitial pneumonia; UIP, usual interstitial pneumonia; IPF, idiopathic pulmonary fibrosis; SD, standard deviation; IQR, interquartile range.

**Supplementary Figure 1**

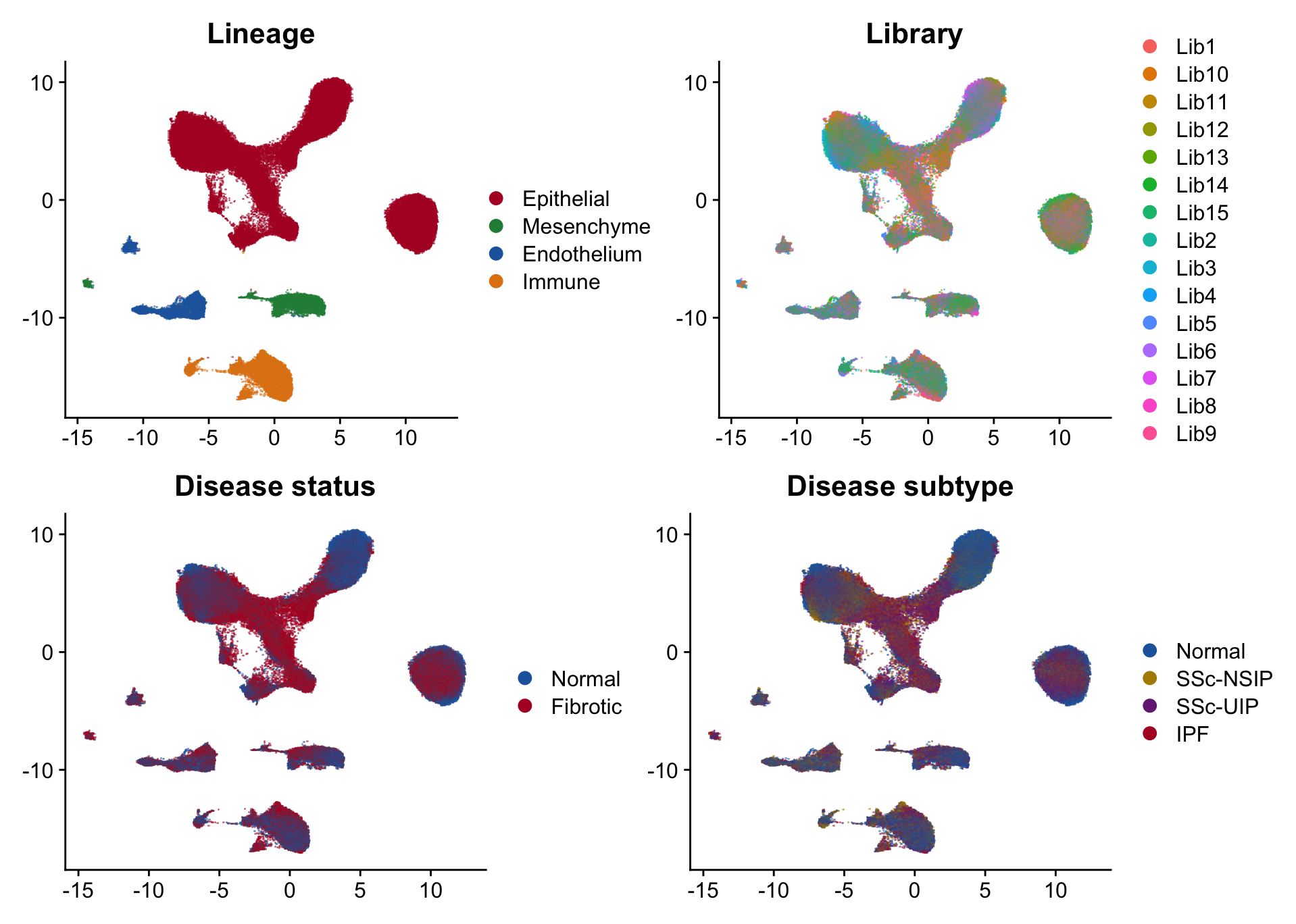

**Supplementary Figure 2**

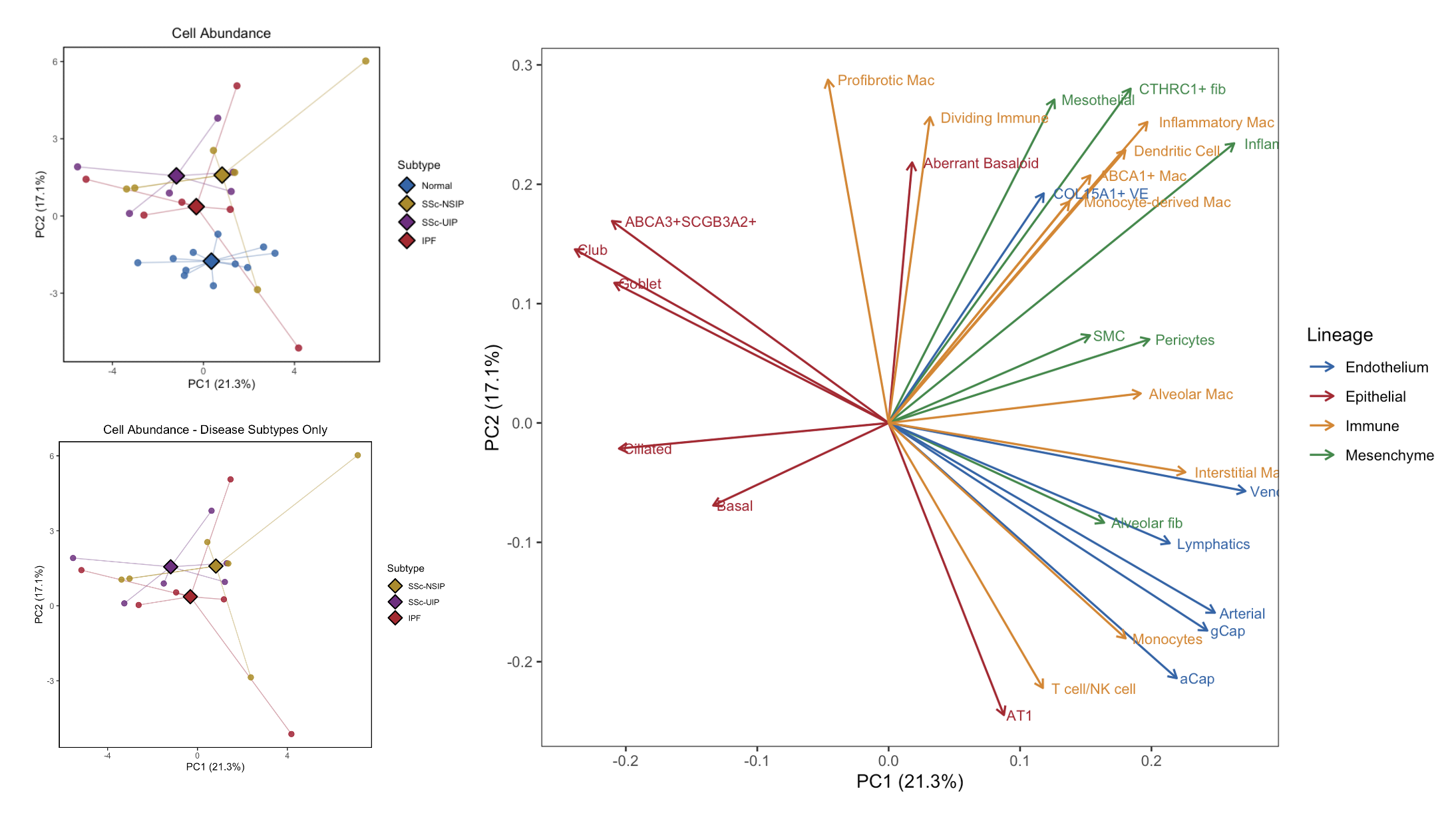

**Supplementary Figure 3**

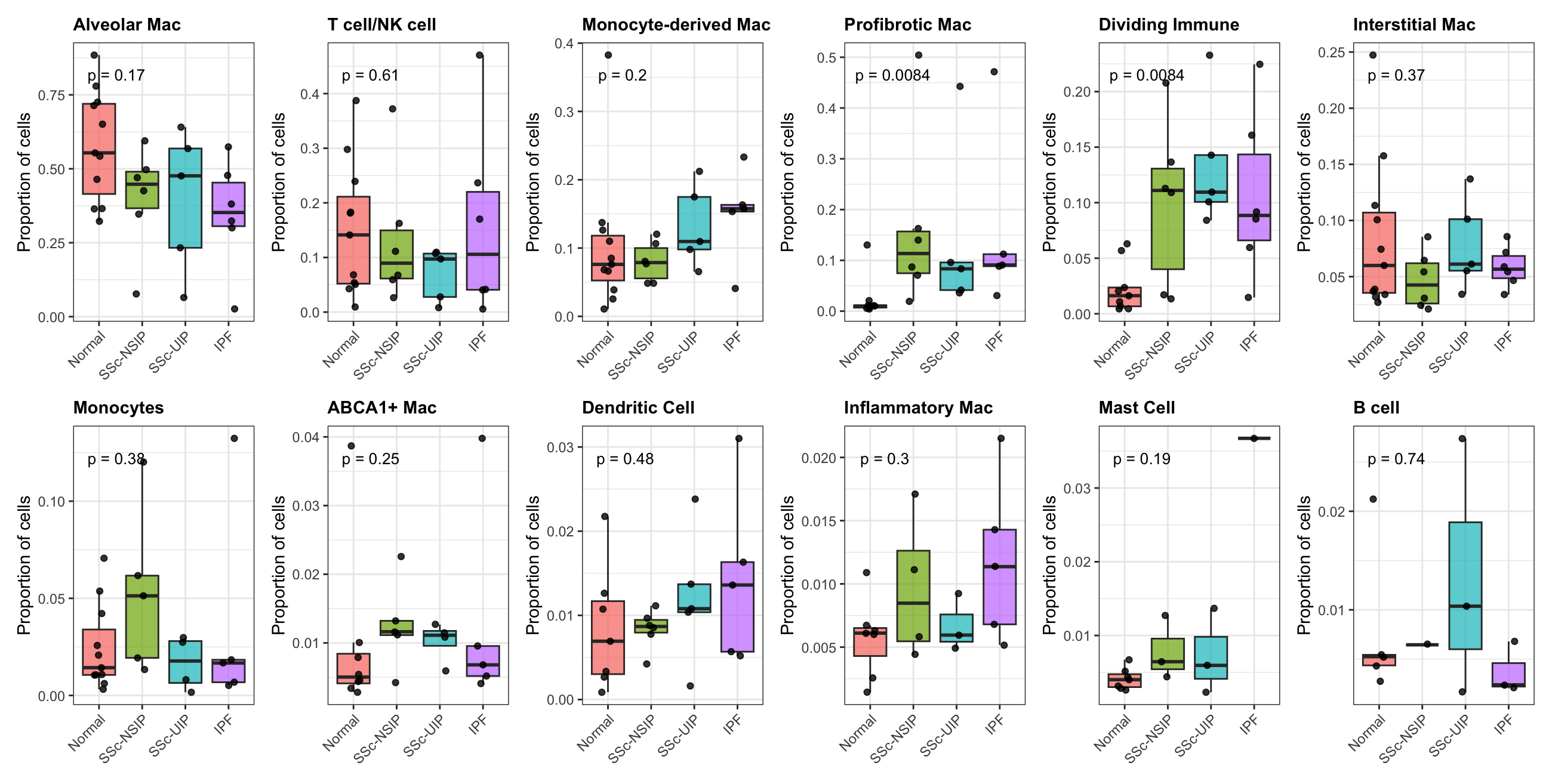

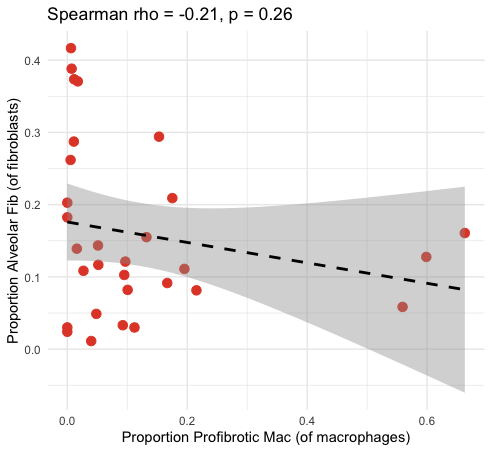

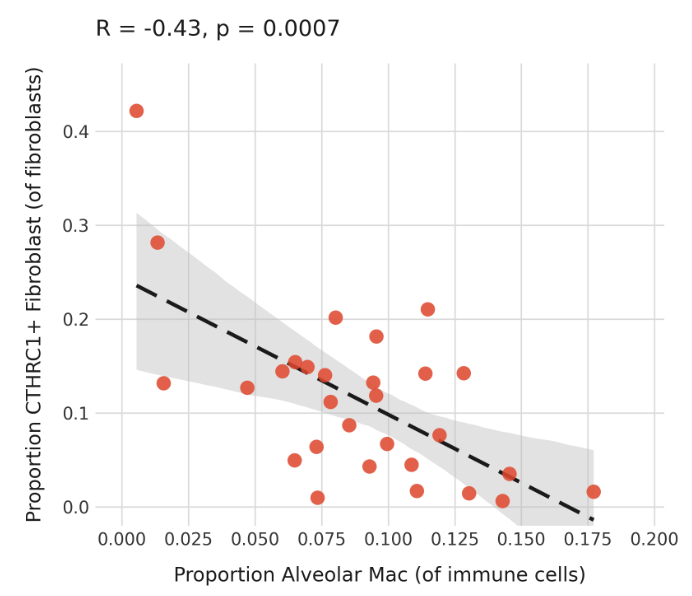
**Supplementary Figure 4:**

**Supplementary Figure 5**

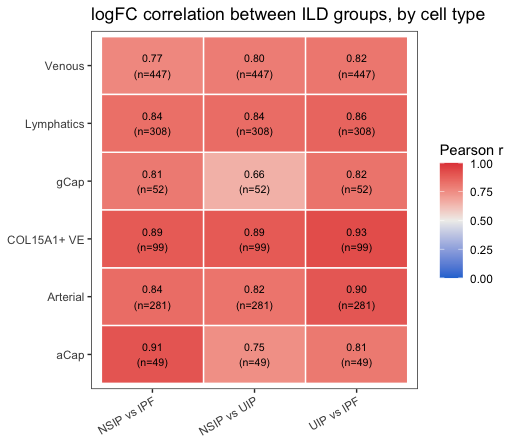

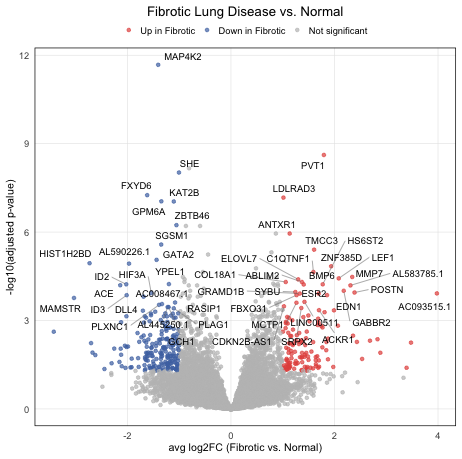

**Supplementary Figure 6**

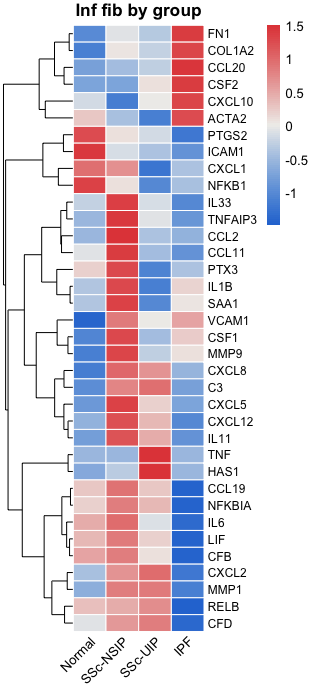

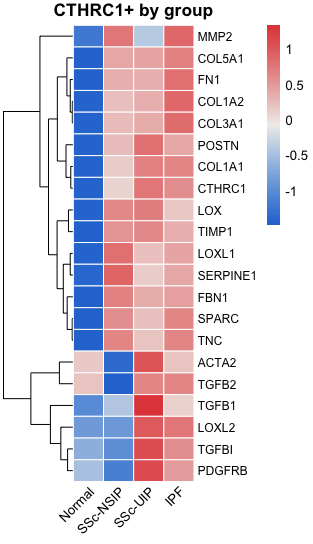

**Supplementary Figure 7**
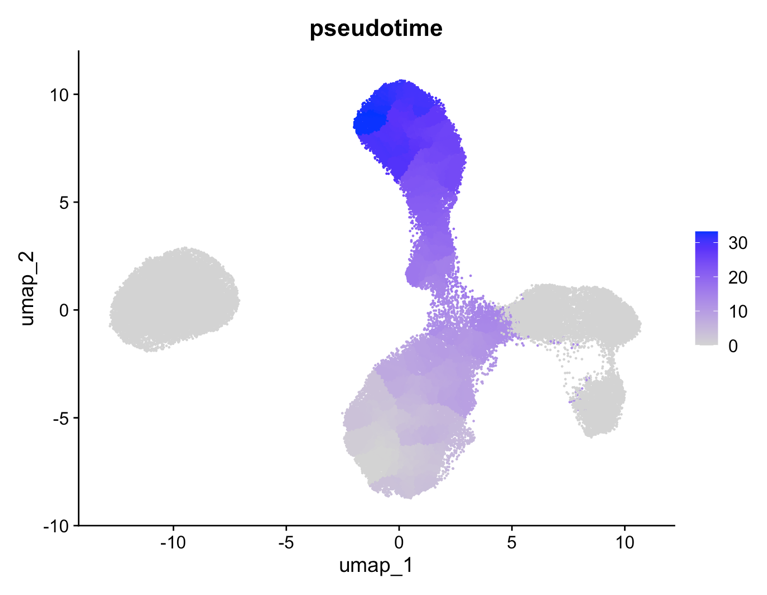

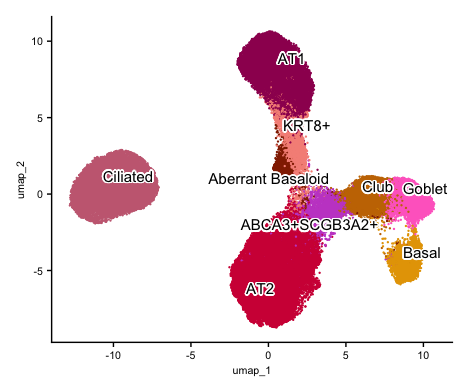

**Supplementary Figure 8**

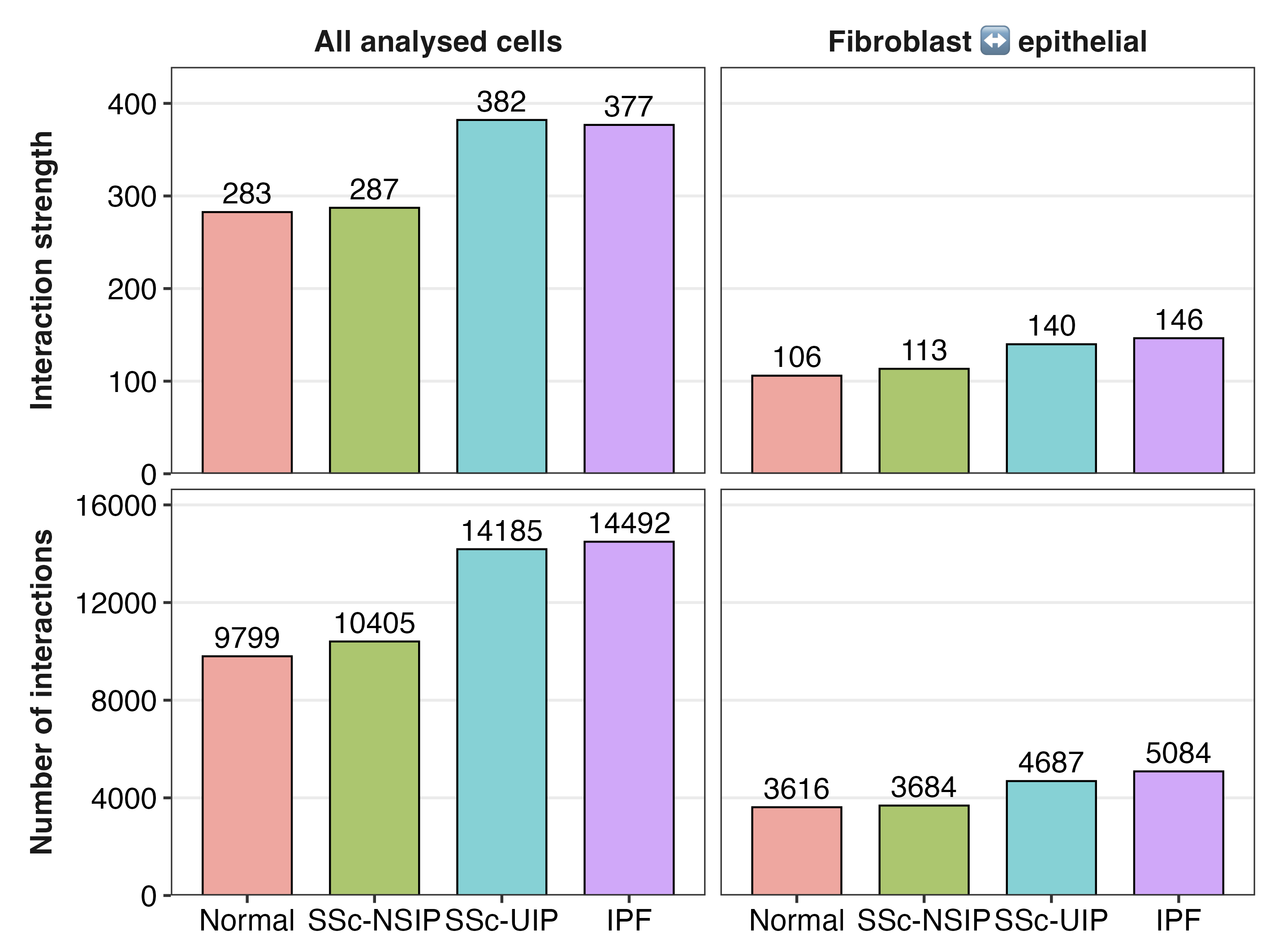

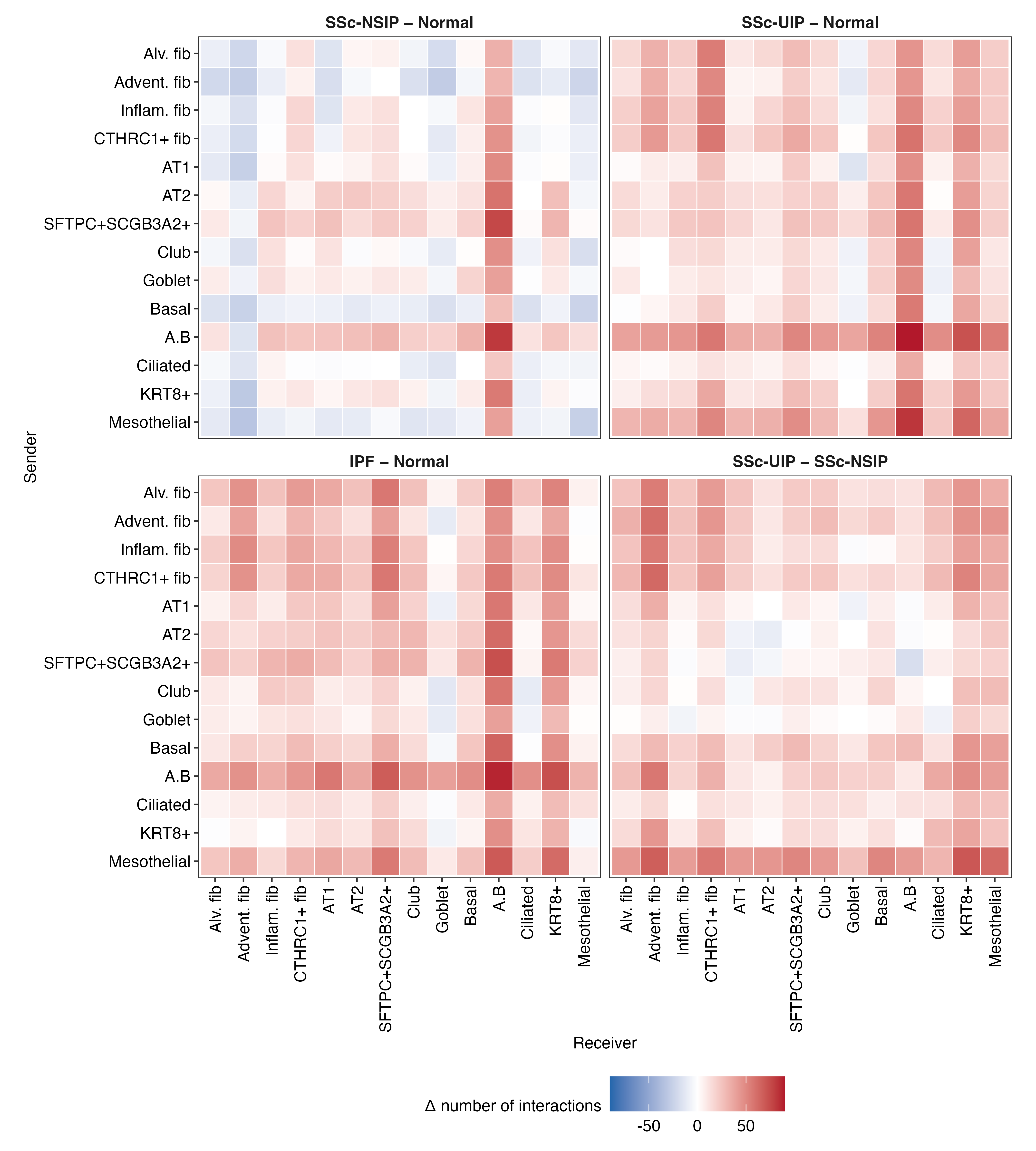

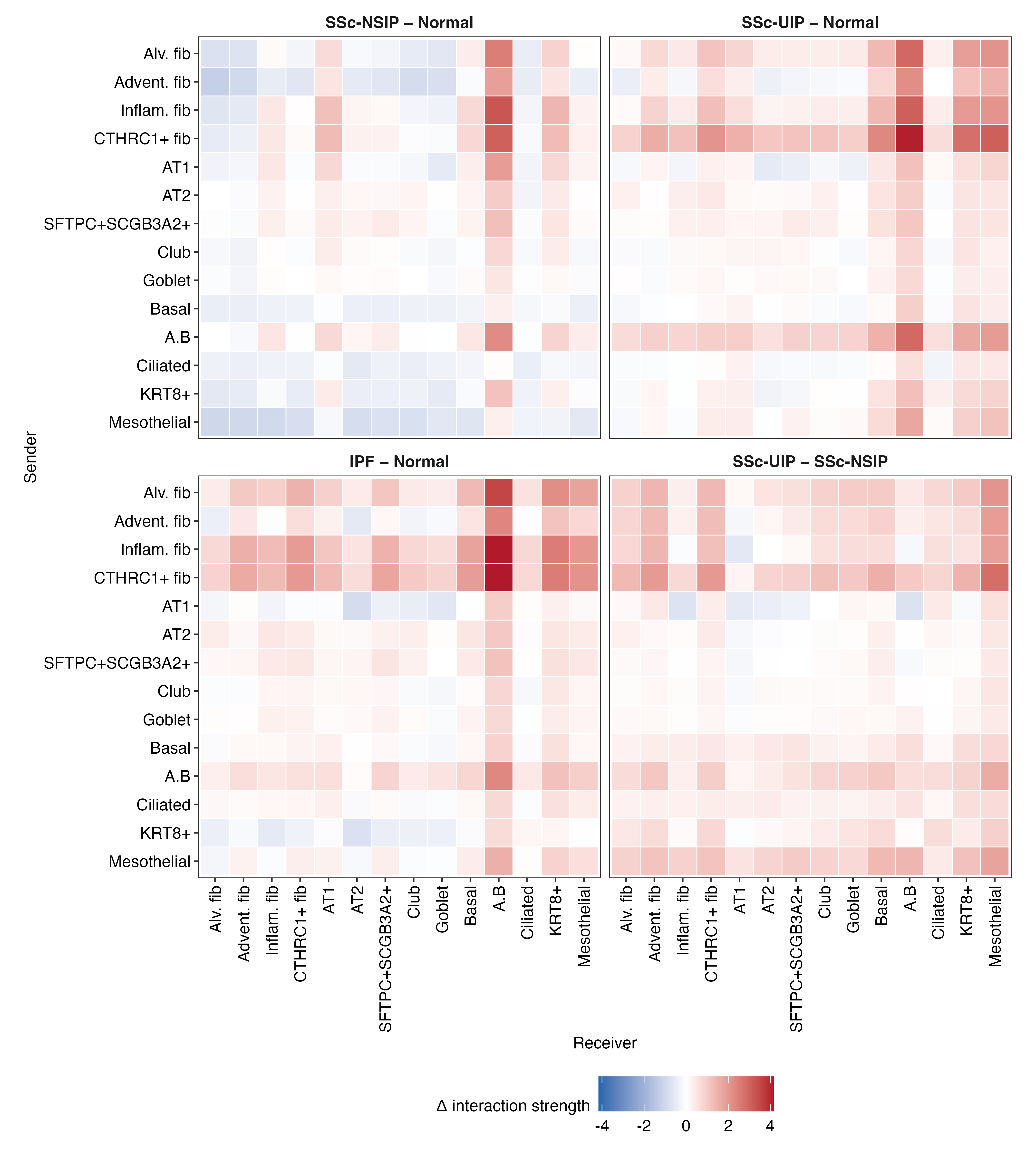
